# Estimating absolute microbial abundances from metabarcoding anchored to cytometry data

**DOI:** 10.64898/2026.07.21.739886

**Authors:** Enrico Ser-Giacomi, Yubin Raut, Jesse McNichol, François Ribalet, Glen Tarran, Christel Hassler, Stephanie Dutkiewicz, Jed A. Fuhrman, Michael J. Follows

## Abstract

Over the past decades, metabarcoding and automated cell-counting approaches have greatly advanced our understanding of marine microbial communities. Metabarcoding provides high taxonomic resolution and comprehensive community characterization, typically as relative gene abundances, whereas flow cytometry provides absolute cell abundances but lower taxonomic coverage. Here, we assess whether concurrent flow-cytometry observations can calibrate metabarcoding data to derive absolute gene abundances across four basin-scale Atlantic and Pacific Ocean transects. We first show that flow-cytometry–anchored calibration reproduces absolute abundances of *Prochlorococcus* and *Synechococcus* with performance comparable (*R*^2^ = 0.87) to internal DNA standard–based quantification. For datasets lacking internal standards, the choice of cytometric “anchor” species introduces systematic offsets in absolute abundance estimates, although spatial patterns remain robust. These offsets may reflect underestimation of cytometric counts or variation in rRNA gene copy numbers among actively dividing cells. We therefore recommend the use of multiple anchors where possible to diagnose systematic uncertainty. Applying this framework, we derive absolute gene concentrations for diverse plankton taxa from compositional metabarcoding data. For taxa with known rRNA gene copy numbers, calibration further enables estimation of absolute cell concentrations. We also resolve ecotype-level absolute abundances of *Prochlorococcus* along a longitudinal temperature gradient, revealing ecological patterns not apparent from compositional or cytometric data alone. Our results demonstrate that calibrated metabarcoding provides a practical quantitative bridge between molecular and cytometric observations, yielding high taxonomic resolution together with absolute gene concentrations and quantified uncertainties.

## 1 Introduction

Diverse microbial communities are a key component of oceanic, freshwater, and soil ecosystems, fueling food webs and mediating global biogeochemical cycles. They are challenging to characterize due to the high abundance of individuals and the wide range of taxonomic resolutions. Despite their importance for understanding global biogeochemical cycles and monitoring environmental change, the biogeography of marine microbes remains only sparsely resolved at the global scale. This is especially true for absolute quantitation, which is rare for ’omics-based analyses of marine microbial community composition. Quantitative characterization of marine microbial populations is achieved using a variety of methods including microscopy, flow cytometry, pigment analysis, remote and in situ optical approaches, qPCR, and metagenomics, each of which has specific advantages and limitations.

Optical cell-count metrics, such as microscopy and flow cytometry, are well established and provide the first benchmark surveys of global-scale marine microbial biogeography (Buitenhuis et al., 2013; Flombaum et al., 2013; Swalwell et al., 2011a; Rees et al., 2017). Cytometric methods typically provide accurate absolute abundance measurements which, combined with some knowledge of cell size, can be re-calibrated as biomass (Ribalet et al., 2019). These data are invaluable for testing and calibrating ecological and biogeochemical simulations (Dutkiewicz et al., 2020; Follett et al., 2022) which, by construction, use mass-based currencies. Optical cell-counting for the whole phytoplankton community requires a combination of instruments to span the large range in cell size (Dutkiewicz et al., 2024) and taxonomic resolution based on morphology has limitations. Notably, most small, non-phototrophic prokaryotes lack any distinctive morphology or pigmentation. Automated imaging platforms, including the Imaging FlowCytoBot (IFCB) and underwater vision profiler (UVP) (Olson and Sosik, 2007a; Picheral et al., 2022), can be leveraged to extend the high throughput potential of monitoring of marine plankton, though unable to taxonomically resolve the smaller pico- and nano-plankton.

Molecular surveys have emerged as a complementary approach. Enabled by technological advances and decreased costs, metagenomic, metatranscriptomic and metabarcoding surveys have emerged as very efficient and powerful approaches for characterizing the composition and structure of microbial communities (Acinas et al., 2021; Biller et al., 2018a; Pesant et al., 2015; Larkin et al., 2021b; Saito et al., 2024; Laiolo et al., 2024). The construction of authoritative reference databases (Quast et al., 2012; Guillou et al., 2012) and development of high-quality primers (Parada et al., 2016; Walters et al., 2016) have further bolstered these approaches. To date, most environmental ’omics surveys have provided compositional data, representing the relative number of specific taxonomic units as a fraction of all the taxonomic units detected in a single sample (dimensionless). Such data have provided new views and valuable insights into marine microbial systems (Sunagawa et al., 2015; Guidi et al., 2016; Sunagawa et al., 2020).

However, absolute abundances (e.g., gene copies l*^−^*^1^) provide more information and significantly extend the power of the data. While most molecular datasets only report relative abundances, a few studies provide absolute cell concentrations inferred from specific, single-copy genes (Pierella Karlusich et al., 2021; Bei et al., 2025). Notably, absolute quantification enables the mapping of spatial or temporal variations in the abundance of a single taxonomic unit, relative to itself (Kurtz et al., 2015). It also facilitates direct inter-calibration and quantitative comparison with other population metrics allowing meaningful compilations of different data types, as well as improving data coverage by enabling the merging of cytometric and genomic observations to create larger, synthesized data sets with greater coverage.

Recent studies have demonstrated close correspondence between relative and absolute abundances of picocyanobacteria from flow cytometry and metabarcoding efforts (Sudek et al., 2015; Jones-Kellett et al., 2024). These findings suggest the potential for a post-calibration of compositional data to absolute abundances (gene copies l*^−^*^1^) for all taxa in molecular datasets where there are concurrent flow cytometric data which can be used to calibrate for a single taxon. Hence, in this study we consider the potential to quantitatively calibrate existing molecular surveys of marine microbes using concurrent cytometric data for a subset of taxa. Our example is focused on calibration of metabarcode data with flow cytometry, though the principles apply to other combinations. Flow cytometry provides absolute abundance data for a limited number of taxa, while metabarcoding provides relative abundance data for a comprehensive community characterization at high taxonomic resolution. If a sample provides cytometric cell concentration for a taxon with a known number of gene copies per cell *and* the relative barcode concentration for that taxon, the cytometric value can calibrate the absolute barcode concentration of that taxon. Furthermore, this calibration can also be applied to all taxa with relative barcode abundances in that sample.

We test and illustrate the intercalibration method using data collected from four basin-scale ocean transects (Figure 1) collected as part of the AMT (Rees et al., 2017), BioGEOTRACES (Biller et al., 2018a), and SCOPE-Gradients (Jones-Kellett et al., 2024) programs. The datasets are described in the Methods section, along with a formal statement of the intercalibration algorithms and associated error propagation. In the Results section, we apply the method and exploit flow cytometric evaluation of *Prochlorococcus* and *Synechococcus* cell concentrations to evaluate absolute environmental barcode abundances for all sequenced taxa in the data sets, revealing biogeographical patterns not discernible from compositional data alone. Exploiting the knowledge of the target gene copy number per cell for both *Prochlorococcus* and *Synechococcus* allows us to test the quantitative skill of this calibration. By leveraging the SCOPE-Gradients metabarcode dataset, which was calibrated with internal biological standards and had concurrent flow cytometry (Jones-Kellett et al., 2024), we also evaluate the errors incurred by intercalibration using flow cytometry alone. Furthermore, we demonstrate the use of multiple cytometric taxa to quantify potential calibration errors in the absence of independent biological standards. In the Discussion section, we examine prospects for obtaining taxonomically resolved, absolute cell abundances for diverse marine plankton with current knowledge of amplification targets per cell (e.g., rRNA copy numbers per cell).

**Figure 1:**
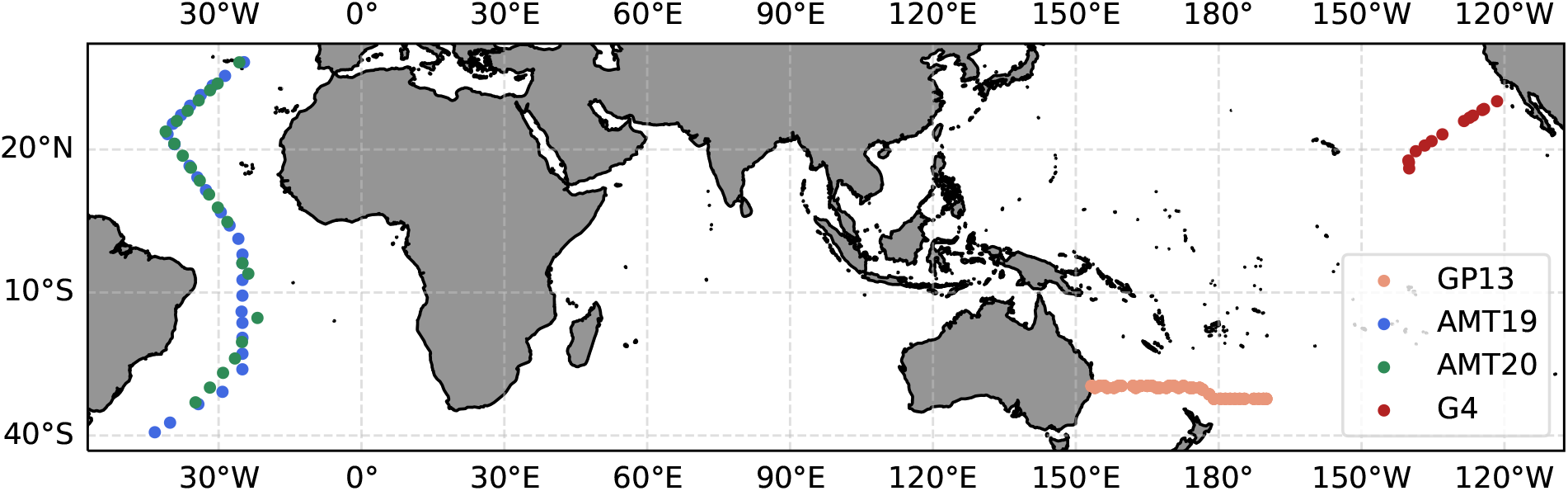
Map of sampling locations of the datasets used in this study: SCOPE-Gradients 4 (Nov. 2021, n = 13, surface only), GP13 (May - June 2011, n = 60, depth-resolved), AMT19 (Oct. - Nov. 2009, n = 28, surface only), and AMT20 (Oct. - Nov. 2010, n = 22, surface only). Each dot represents the sampling site and colors indicate the corresponding dataset.

## 2 Materials and Procedures

### 2.1 Datasets

Metabarcoding is high throughput, efficiently providing significant data coverage, and provides a high level of taxonomic resolution. It can provide a very comprehensive characterization of the microbial community, depending on the primer set (McNichol et al., 2021). We leveraged four whole-community Amplicon Sequence Variant (ASV) datasets, generated with the 515Y/926R primers (Parada et al., 2016), paired with high-quality flow cytometric measurements of picoplankton population concentrations. Two of these datasets represent surface samples collected in 2009/2010 during the AMT-19/20 cruises (an ongoing sampling campaign from England to Tierra del Fuego). The third dataset is a depth-resolved transect collected in 2011 during the first leg of the GEOTRACES cruise GP13 from Australia to New Zealand. All ASVs from these three transects (single sample replicates) are part of the Global rRNA Universal Metabarcoding of Plankton (GRUMP) dataset (v1.3.5) compilation (https://github.com/jcmcnch/Global-rRNA-Univeral-Metabarcoding-of-Plankton)(McNichol et al., 2025). Flow cytometric data (single sample replicates) from the AMT cruises was obtained from the Simons Collaborative Marine Atlas Project (CMAP) website (Ashkezari et al., 2021), and for GP13 was generated by Christel Hassler (Cabanes et al., 2020). A fourth dataset was collected in 2021 in the Eastern North Pacific as part of the SCOPE-Gradients G4-cruise (Jones-Kellett et al., 2024). This data set included replicate metabarcoding, calibrated by internal biological standards, samples as well as concurrent flow cytometry (BD Influx) measurements in triplicates.

The ASV datasets generated from these samples had two important properties that made them suitable for this study: DNA extractions were carried out on unfractionated samples (*>*0.2 *µm*) and ASVs were generated using a primer pair that is free of any major primer mismatches for taxa found in oxic seawater (McNichol et al., 2025). Consequently, the resulting data should be consistent and free of major biases (Parada et al., 2016; Yeh et al., 2021; McNichol et al., 2021). The primer pair perfectly matches 96% of all rRNA sequences found in a global-scale, marine metagenomic dataset, and amplicon abundances generated from these primers accurately reflected those obtained from metagenomes when tested on two long transects (McNichol et al., 2021).

### 2.2 Metabarcoding-based methods background

For all metabarcode-based methods, a target gene (e.g., SSU rRNA) is used to classify and quantify the abundance of individual taxa. Quantification of this target gene can be determined with amplification-based approaches (i.e., PCR) or amplification-free techniques (i.e., shotgun metagenomics). After DNA sequencing of many tens of thousands of barcodes per sample, barcodes are processed, often using Operational Taxonomic Unit (OTU) clustering or “denoising” to Amplicon Sequence Variants (ASV), to remove sequencing errors and merge similar or identical sequences (Blaxter et al., 2005; Callahan et al., 2016, 2017). Each of these merged barcodes are then taxonomically classified by comparison with a reference database. The final result is a count table where each unit (OTU or ASV) is associated with a taxonomy and count data for each individual sample.

### 2.3 Intercalibration algorithm and uncertainties

Here we formalize a general framework, described conceptually in the Introduction, which relates the relative number of taxon-specific, raw metabarcoding reads derived from DNA sequencing to absolute environmental barcode concentrations by leveraging independent cell count data of an “anchor” taxon (e.g., from flow cytometry or any other cell counting methods). We consider a generic set of independent samples labeled hereafter with the superscript *j* and refer to a specific taxonomic assignment using the subscript *i*. A sample *j* represents a single, independent environmental observation from which we obtain a characterization of the local microbial community. We intentionally do not provide a genetic/phylogenetic definition of taxon, *i* as this framework is agnostic and can be used at a highly resolved level (e.g., *Prochlorococcus* ecotype) or at broad functional level (e.g., picoeukaryote), so long as the investigator can define their categories of choice confidently from metabarcode taxonomic assignments. However, we note that in practice, the broader the level, the more unlikely it is that it will have uniform characteristics.

Metabarcode data are most often presented in relative terms, where *R_i_^j^* is the fraction of total reads associated with taxon *i* in sample *j*. Table 1 defines symbols and units referenced in the following equations. *R_i_^j^* is related to the associated environmental gene concentration, *k_i_^j^* as follows:

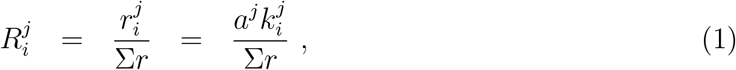

**Table 1:** Symbols, names, and dimensions of the model variables.

| Variable | Name | Units |
| --- | --- | --- |
| $r_i^j$ | Metabarcode sequencing reads associated with taxon $i$ in sample $j$ | $\# \text{ reads}$ |
| $\sum_{i=1}^n r_i^j$ | Sum of metabarcode reads from $n$ selected taxa, $i$, in sample $j$ | $\# \text{ reads}$ |
| $R_i^j = \frac{r_i^j}{\sum_{i=1}^n r_i^j}$ | Fraction of reads associated with taxon $i$ in sample $j$ | non-dimensional |
| $k_i^j$ | Gene copies per volume associated with taxon $i$ in sample $j$ | $\frac{\# \text{ copies}}{\text{volume}}$ |
| $t_i$ | Number of target gene copies per cell for taxon $i$ | $\frac{\# \text{ copies}}{\text{cell}}$ |
| $a^j$ | Sample-specific multiplication factor for sample $j$ | $\frac{\text{volume} \# \text{ reads}}{\# \text{ copies}}$ |
| $n_i^j$ | Cell counts per volume of taxon $i$ in sample $j$ | $\frac{\# \text{ cells}}{\text{volume}}$ |

where the normalizing factor, Σ*r* = Σ*_i_*_=1_^*n*^*r_i_^j^* is the sum of all reads in the sample, or the sum of selected reference subset. The sample-specific multiplication factor, *a_j_*, is the ratio of metabarcoding reads to the number of total amplification targets associated with taxon *i* present in sample *j*. The multiplication factor *a^j^*, is assumed independent of *i*; in other words, there are no taxon-specific amplification biases that would result in the preferential amplification of one taxon over another. The representativeness of relative abundances also depends upon this assumption.

The relative barcode abundance, *R_i_^j^*, can also be related to the cell concentration, *n_i_^j^*, in the environmental sample by accounting for the number of target gene copies per cell *t_i_*:

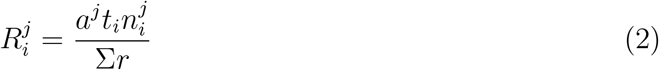

This relationship can be useful for testing, calibration and merging of data sets in conjunction with flow cytometry or microscope data, for example. The number of target gene copies per cell *t_i_* may be derived from expert knowledge of a given organism. For the case of SSU rRNA metabarcoding, *t_i_* represents an operational measure of the number of targeted genes per cell. For targeted, amplicon-based approaches (i.e., a 16S- or 18S-specific primer set) *t_i_* is defined as the number of 16S or 18S copies per cell. For PCR primers that can amplify both 16S and 18S SSU rRNA from a eukaryotic phytoplankter, *t_i_* can be defined as the number of copies of one or both of these target genes (Yeh et al., 2021). Equation (2) assumes that the number of amplification targets per cell *t_i_* is sample independent and a conserved “trait” of each taxon *i*. This is thought to be a good approximation for many prokaryotic organisms (Gao and Wu, 2023; Martin et al., 2022) but may be less appropriate for others including dinoflagellates (Ruvindy et al., 2023).

### 2.4 Estimating absolute environmental gene copy concentration and cell concentration

In this study, we ask: how accurately we can calibrate absolute environmental gene concentrations, *k_i_^j^*, from relative metabarcode reads, *R_i_^j^*, by exploiting cell counts, *n_i_^j^*, from concurrent flow cytometry? To do so, we first evaluate the multiplication factors, *a^j^*, by exploiting an *anchor taxon*, denoted by subscript “*c*”. The anchor taxon must be quantified with metabarcoding reads (*R_c_^j^*) as well as cell counts (*n_c_^j^*) from flow cytometry (or microscopy, CARD-FISH, etc.) associated with all samples. Its amplification targets per cell (*t_c_*) must also be known. Rearranging Eq. (2), we can then estimate the unique multiplication factor for each sample, *a^j^*:

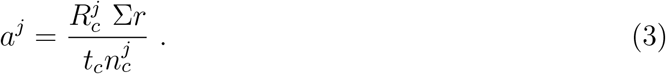

Then we can evaluate the absolute environmental gene copy concentration, *k_i_^j^* for all taxa using Eq. 1 and the quantified amplification factors, *a^j^*:

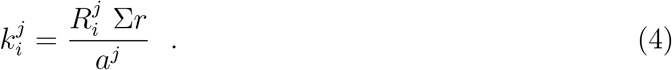

Here *k_i_^j^* is an absolute measure of taxonomic abundance (gene copies per unit volume) that allows quantitative comparison both within and across samples (Lin et al., 2019). Note that this derivation of *k_i_^j^* does not rely on *a priori* knowledge of barcode copy numbers per cell (i.e., *t_i_*) other than that of the anchor taxon. It provides an absolute quantification of barcode-derived environmental gene concentration that is, in theory, identical to that derived from the internal nucleic acid standard approach.

For any taxon for which we do have knowledge of *t_i_* we can also evaluate the absolute environmental cell concentration (cells m*^−^*^3^) by applying Eq. 2:

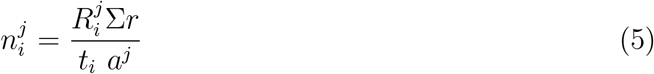

providing a direct interface between cytometric and metabarcode quantities.

### 2.5 Evaluating Uncertainties

Importantly, all the input data have associated uncertainties (Δ) which can be quantified and we can propagate the errors to provide uncertainties on the absolute environmental gene and cell concentrations. For example, Δ*n_i_^j^*, the uncertainty associated with the estimation of the absolute cell concentration of taxon-*i* in sample-*j* can be estimated using standard first-order error propagation (Taylor, 2022).

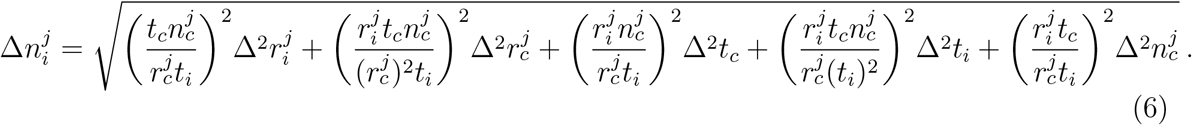

Where Δ*t_c_,* Δ*t_i_,* Δ*r_i_,* Δ*r_c_,* Δ*n_c_* are uncertainties over *t_c_, t_i_, r_i_, r_c_, n_c_*. An equivalent expression can be defined for absolute environmental gene concentrations, *k_i_^j^*.

#### 2.5.1 Quantifying uncertainties for input data

Cell concentrations evaluated by flow cytometry are subject to uncertainty from several sources, including weak fluorescence of picocyanobacteria acclimated to high light intensity, such as *Prochlorococcus* (Phongphattarawat et al., 2023), and limitations of instrument performance (Gérikas Ribeiro et al., 2016). Comparing two flow cytometers from the same manufacturer, one sufficiently portable for use in the field (BD Accuri C6) and the other less so (BD FACSCanto), Gérikas Ribeiro et al. (2016) reported an approximately 25% inter-instrument difference in estimated *Prochlorococcus* and *Synechococcus* cell concentrations from the same samples collected in the Tropical Atlantic euphotic zone. Instrumental differences, and the associated uncertainties, were considerably lower for larger nanophytoplankton, which carry a larger pigment quota (Gérikas Ribeiro et al., 2016). A recent study examined triplicate flow cytometric evaluations of surface *Prochlorococcus* and *Synechococcus* cell concentrations in the North Pacific (Jones-Kellett et al., 2024), finding 11% and 15% errors respectively relative to the sample means for more than 95% of the samples. Informed by these studies, we assume conservative technical uncertainties, Δ*n_c_^j^*, of 15% for flow cytometric picocyanobacteria concentrations used in this work, where most of the samples were obtained from near-surface, high-light waters.

Metabarcode-based estimates of relative abundance, *R_i_^j^*, and the underlying quantification of reads, *r_i_^j^*, have implicit methodological uncertainties. Triplicate analyses of metabarcode abundances from single samples, made using the same protocols as in this study (Jones-Kellett et al., 2024), suggest an uncertainty on individual relative metabarcode abundances, Δ*r_i_^j^*, of approximately 15% of the sample mean for *Prochlorococcus* and *Synechococcus*. Here we assume this value for all taxa.

The variation of DNA content over the cell cycle (Vaulot, 1995) leads to a potential time- and taxon-dependent source of uncertainty in *t_i_* which we acknowledge but have not accounted for here.

## 3 Assessment

We analyzed planktonic communities sampled during four independent oceanographic campaigns: GP13 (BioGEOTRACES), AMT19 and AMT20 (Atlantic Meridional Transect program), and G4 (SCOPE-Gradients) (see Fig. 1). For each transect, concurrent metabarcoding and flow cytometric cell counts are available (Methods). In this section, we demonstrate several applications of the absolute abundance calibration framework to:

1. estimate absolute *Prochlorococcus* and *Synechococcus* cell concentrations from relative metabarcode data with associated uncertainties and validate them against parallel flow cytometric cell counts and quantitative abundances derived from amplicon sequencing with internal genomic standards;
2. evaluate the range of *a_j_* ’s and sensitivity of the intercalibration approach to the choice of the anchor taxon; and
3. provide estimates for additional non-cyanobacterial taxa in samples with parallel metabarcoding and enumeration data obtained from flow cytometry along the AMT-19, AMT-20, and GP13 transects.

### 3.1 Quantitative agreement of *Prochlorococcus* and *Synechococcus* cell abundances between calibrated ASV-based estimates and enumeration by flow cytometry and internal standards

Our desire is to use cell concentrations from flow cytometry to calibrate relative gene concentrations, obtained from metabarcodes, into absolute gene concentrations. While the principle is straightforward, expressed in the algorithms in the Methods section, merging data types in this way will also introduce additional uncertainties. In order to address the potential for this approach, we leverage an existing data set for which relative gene concentrations, absolute gene concentrations calibrated with internal biological standards, and flow cytometry were all available (Jones-Kellett et al., 2024). This data set allows us to directly compare estimated absolute cell concentrations for *Prochlorococcus* and *Synechococcus* derived from metabarcode data calibrated by genetic standards and when calibrated by flow cytometry.

In that data set, Jones-Kellett et al. (2024) showed that the absolute cell concentrations recovered from metabarcode data calibrated with internal genomic standards, and assuming the well-established values of *t_i_* for these taxa, match extremely well with independent flow cytometric evaluations (see Figure 2(a), which recapitulates their finding). As a proof of concept, we first asked how well the absolute cell concentrations would have been recovered from the metabarcode data had we, instead of using biological standards, calibrated *Prochlorococcus* cell concentrations anchored by flow cytometry for *Synechococcus* and vice versa.

**Figure 2:**
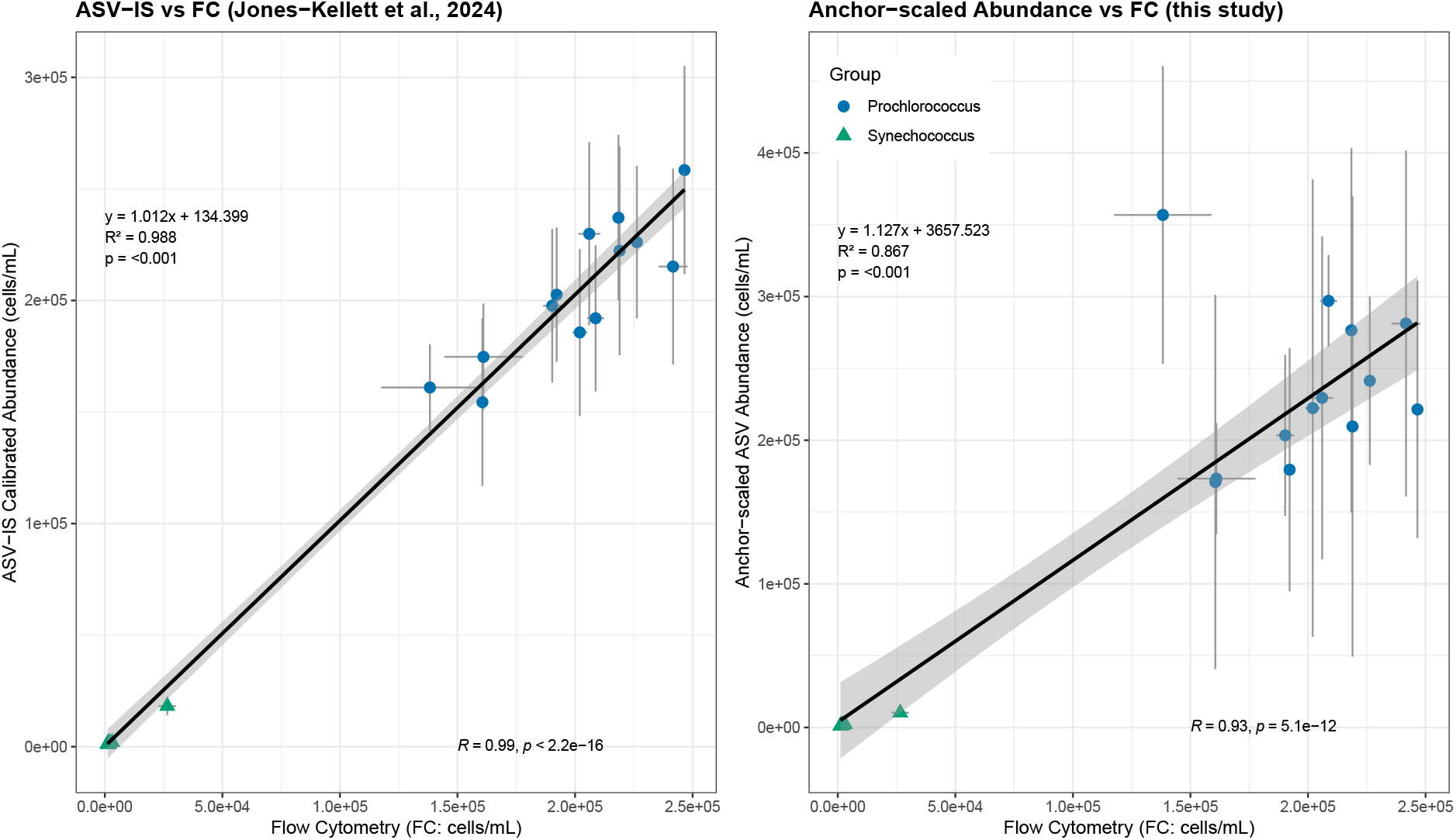
(a) Absolute cell concentration (cells mL*^−^*^1^) of *Prochlorococcus* and *Synechococcus* from the G4 transect (n = 13) derived from metabarcode data using internal DNA standards (ASV-IS calibrated abundance), shown against flow cytometric cell concentrations (Jones-Kellett et al., 2024). (b) Anchor-scaled ASV abundance of *Prochloroccus* and *Synechococcus* from the same samples (n = 13), obtained by calibrating ASV-derived abundances using flow-cytometric *Synechococcus* and *Prochloroccus* as anchor taxa, shown against flow cytometric cell concentrations (this study).

First, we rescaled the compositional metabarcode data, *R_i_^j^*, into absolute environmental gene concentrations, *k_i_^j^*, using (Eq. 4). To do so, we evaluated the sample-specific multiplication factor (*a^j^*; Eq. 3) for each sample *j* based on concurrent flow cytometric cell counts and metabarcode reads of an anchor taxon for each sample along the Gradients-4 transect. Given their well-established *t_i_* values, *t*_pro_ = 1 and *t*_syn_ = 2 (Dufresne et al., 2003; Fuller et al., 2003; Schirrmeister et al., 2012; Stoddard et al., 2015), we then estimated absolute cell concentrations using Eq. 5, providing a direct calibration between ASV-based relative read counts and cytometric cell enumerations.

*Synechococcus* was used as the anchor to evaluate *a^j^*, and a calibrated, ASV-based estimate of absolute cell concentration for *Prochlorococcus* and vice versa. Across the G4 transect, these cytometrically-calibrated, ASV-based cell abundances for both *Prochlorococcus* and *Synechococcus* closely matched those obtained by direct flow-cytometry as well as those obtained using internal genomic standards (Fig. 2). When using flow cytometric counts as the anchor, ASV-derived estimates reproduced a near 1:1 relationship for both taxa (Fig. 2b; slope of 1.13 and Pearson *r* = 0.93), mirroring the strong quantitative agreement previously observed between internal-standard–derived and cytometric measurements (Fig. 2a; slope = 1.01 and Pearson *r* = 0.99; Jones-Kellett et al. (2024)). This correspondence demonstrates that the calibration framework can recover absolute abundances from compositional metabarcoding data with high fidelity.

Error propagation analysis (Eq. 6) revealed that uncertainty in ASV-based cell estimates scales directly with the variability in the anchor taxon’s enumeration. Samples exhibiting greater dispersion in flow cytometric counts of the anchor taxon displayed correspondingly larger propagated uncertainties in the calibrated estimates (Supplementary Fig. S1). This highlights the importance of assessing the stability and precision of the anchor measurements prior to calibration, as uncertainty in anchor quantification can amplify downstream errors in derived abundances.

### 3.2 Calibrating with flow cytometry: Senstivity to anchor taxon

Having established that the calibration framework can quantitatively and accurately reproduce absolute cell abundances when all key components are available, we next evaluated its robustness when applied to historical datasets lacking internal genomic standards. Specifically, we used cross-calibration between the major picocyanobacteria taxa *Prochlorococcus* and *Synechococcus* sampled across the Atlantic Ocean (AMT-19, AMT-20) and South Pacific Ocean (GP13) to assess the consistency of sample-specific multiplication factors and to detect potential systematic biases.

Using Eq. (3) with either *Prochlorococcus* or *Synechococcus* as the anchor taxon, we evaluated the multiplication factors, *a_j_*, for all samples across these three transects. Panel A in Figure 3 shows the distribution of sample-specific multiplication factors (*a^j^*) calculated using *Prochlorococcus* as the anchor taxon. This reveals that *a^j^* values vary by roughly an order of magnitude within a single transect and can differ among cruises, reflecting variability in sequencing depth and sample-specific amplification efficiency. Panel B compares *a^j^* values derived from *Synechococcus* versus *Prochlorococcus* as anchors which, in theory, should be identical. In this case we find that *Synechococcus*-based *a^j^* are systematically lower than *Prochlorococcus*-based values. Linear regression (slope = 0.321, intercept = 0.027, *R*^2^ = 0.22) indicates a weak but significant correlation (Pearson *r* = 0.47, *p* = 5 × 10*^−^*^7^). Deviations from the 1:1 line reflect sample- or taxon-specific biases in amplification or cell counts.

**Figure 3:**
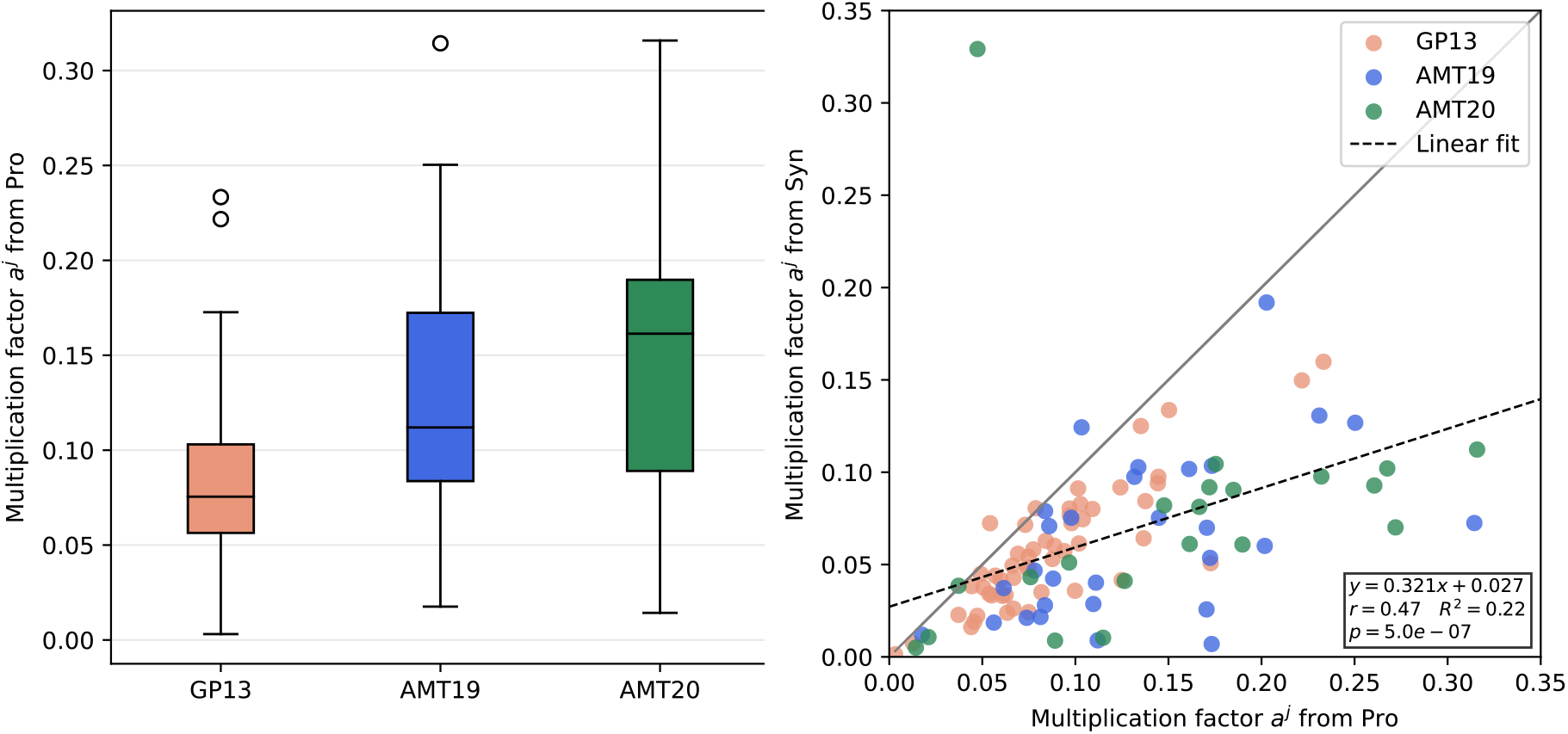
(a) Boxplots of the *a_j,_*_pro_ distributions for the three datasets analyzed. Each boxplot is associated with a single transect. (b) Relationship of the corresponding *a_j_* values derived using *Prochlorococcus* and *Synechococcus* anchors respectively.

The anchor-dependence of *a_j_* propagates through to absolute abundance estimates. We applied the cross-anchoring approach, using *Synechococcus* as the anchor to estimate *Prochlorococcus* abundances and vice versa (Figure 4). Across the AMT-19, AMT-20, and GP13 transects, calibrated ASV-based estimates of *Prochlorococcus* and *Synechococcus* cell concentrations were highly correlated with direct flow-cytometric measurements over four orders of magnitude (Fig. 4). ASV-calibrated estimates of *Prochlorococcus* cell concentrations using *Synechococcus* as the anchor were generally higher than the corresponding flow-cytometry counts (Fig. 4A; Pearson r = 0.82, p-value *<* 0.001), reflecting the systematic differences in the estimates of *a_j_*. Conversely, ASV-calibrated *Synechococcus* cell concentrations using *Prochlorococcus* as the anchor were generally lower than flow-cytometry counts, with most points below the 1:1 line, while also preserving strong positive correlation (Fig. 4B; Pearson r = 0.90, p-value *<* 0.001). These reciprocal systematic shifts are consistent across all three transects and align with the observed differences in *a_j_* values derived from the two anchors (Fig. 3).

**Figure 4:**
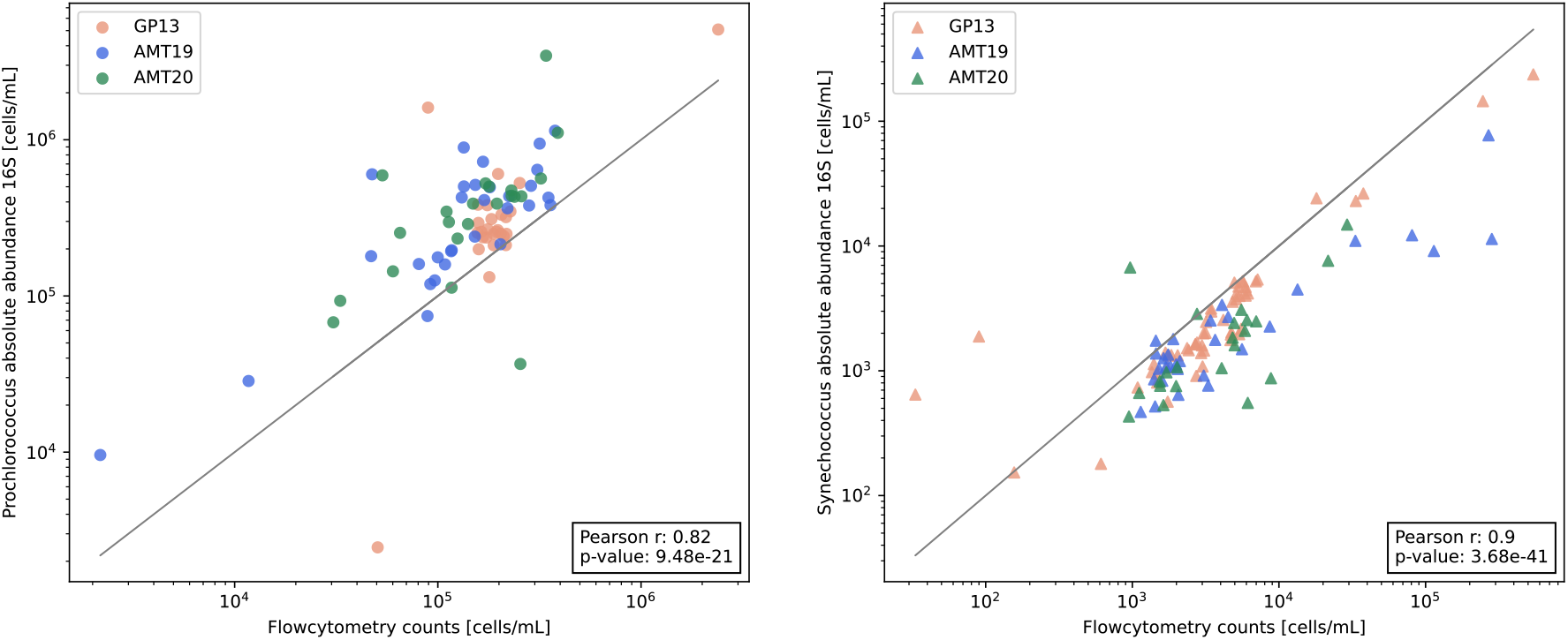
(a) Scatter plot of absolute cell abundances of *Prochlorococcus* inferred from ASV data (y-axis), using *Synechococcus* as the anchor taxon, versus measurements from flow cytometry (x-axis). (b) Scatter plot of absolute cell abundances of *Synechococcus* inferred from ASV data (y-axis), using *Prochlorococcus* as the anchor taxon, versus measurements from flow cytometry (x-axis). Each data point corresponds to a single sampling site, different colors are associated with different datasets (GP13 - orange, AMT19 - blue, AMT20 - green).

This systematic difference is consistent with documented challenges in characterizing dim *Prochlorococcus* populations in high-light surface waters (Gérikas Ribeiro et al., 2016; Phongphattarawat et al., 2023), where most of these samples were collected, potentially resulting in underestimation of *Prochlorococcus* by flow cytometric enumeration. The variability of cell concentrations of *Synechococcus*, which can be very low, may also introduce higher uncertainties or potentially inflate *Prochlorococcus* estimates for some samples when used as an anchor (Supplemental Figure S1). Additionally, this variability can also be driven by some of the cells being in mitosis, which could potentially fluctuate *t_i_* values (Vaulot, 1995).

Together, these results demonstrate that cross-anchoring preserves relative abundance patterns but can also reveal potential systematic offsets, revealing additional potential uncertainties, or careful choice of the more reliable anchor under a particular given circumstance. This highlights the diagnostic value of using multiple anchor taxa for calibration.

### 3.3 Extending ASV-calibration to non-cyanobacteria taxa

Using Eq. (3) with *Synechococcus* as the anchor taxon, we evaluated the multiplication factors, *a_j_* for all samples across three independent transects: AMT-19, AMT-20, and GP13. With these empirically derived *a^j^*values, we estimated the absolute environmental gene copy concentrations (*k_i_^j^*) for all taxa in each sample (Eq. 4). This transformation converts relative sequence abundances into directly comparable quantitative metrics, without requiring prior knowledge of barcode copy numbers per cell (*t_i_*).

As an illustration, a broad comparison of the 15 most abundant taxa along the AMT-19 transect highlights that community structure inferred from relative abundances can differ markedly from that based on absolute gene concentrations (Fig. 5). The cumulative contribution of the fifteen most abundant taxa exhibit different trends (e.g., 10°S - 20°N) between relative abundance and absolute gene concentration (Fig. 5). Moreover, individual taxa (e.g., SAR11-Ia: ASV4, SAR11-II: ASV7) also show differing patterns of relative abundance and gene concentration across samples throughout the transect (Supplementary Fig. S2). This decoupling highlights that compositional data alone can obscure real variation in total gene content and community structure along the transect, whereas ASV-calibrated gene concentrations capture meaningful changes in abundance across sites.

**Figure 5:**
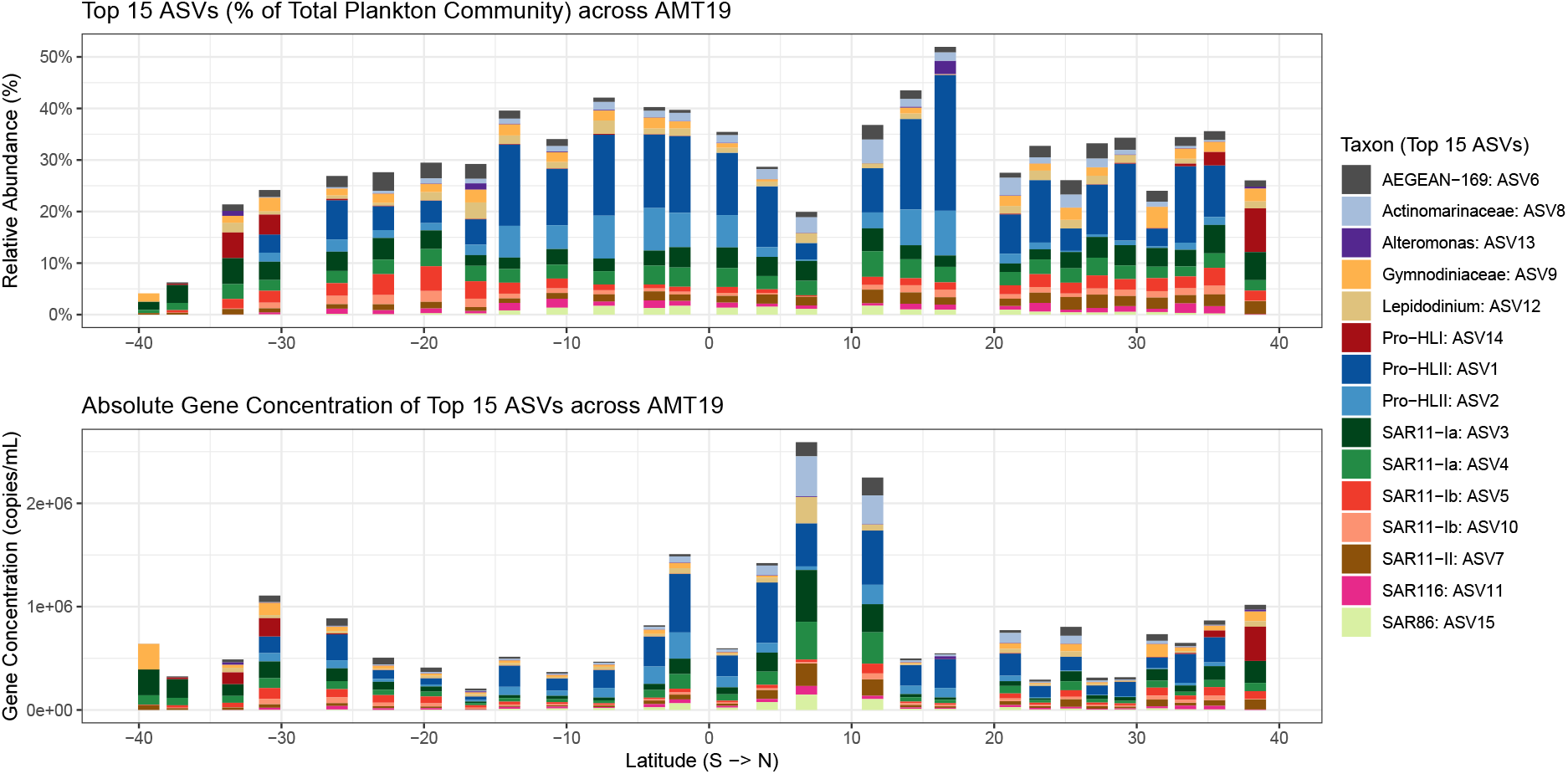
Relative (a) and absolute (b) environmental gene concentration of the fifteen most abundant taxa across the AMT19 transect (n = 28). Absolute concentrations were calibrated using *Synechococcus* anchor. The gene concentration estimates from using *Prochlorococcus* as an anchor and the comparison with *Syn*-anchored estimates are highlighted in (Supplementary Figs. S2, S3).

For a subset of plankton taxa (e.g., picoeukaryotes, nanophytoplankton, dinoflagellates, ciliates) with documented *t_i_* values derived largely from culture studies, we further converted gene concentrations into cell concentrations using their specific *t_i_* values (Martin et al., 2022). Fig. 6 compares relative abundances and estimated cell concentrations for these taxa across all representative samples from the 3 transects with these taxa present. Across samples, these conversions reveal systematic shifts compared to relative abundance metrics. Larger organisms (e.g., nanophytoplankton, dinoflagellates) with higher rRNA copy numbers contribute in a similar range of relative abundance as other samples dominated by picoeukaryotic plankton with lower rRNA copy numbers. However, this over representation in relative abundance is corrected for when calibrated to cell concentrations which show that cell counts for the larger plankton are often orders of magnitude lower than picoeukaryotic phytoplankton (Fig. 6).

**Figure 6:**
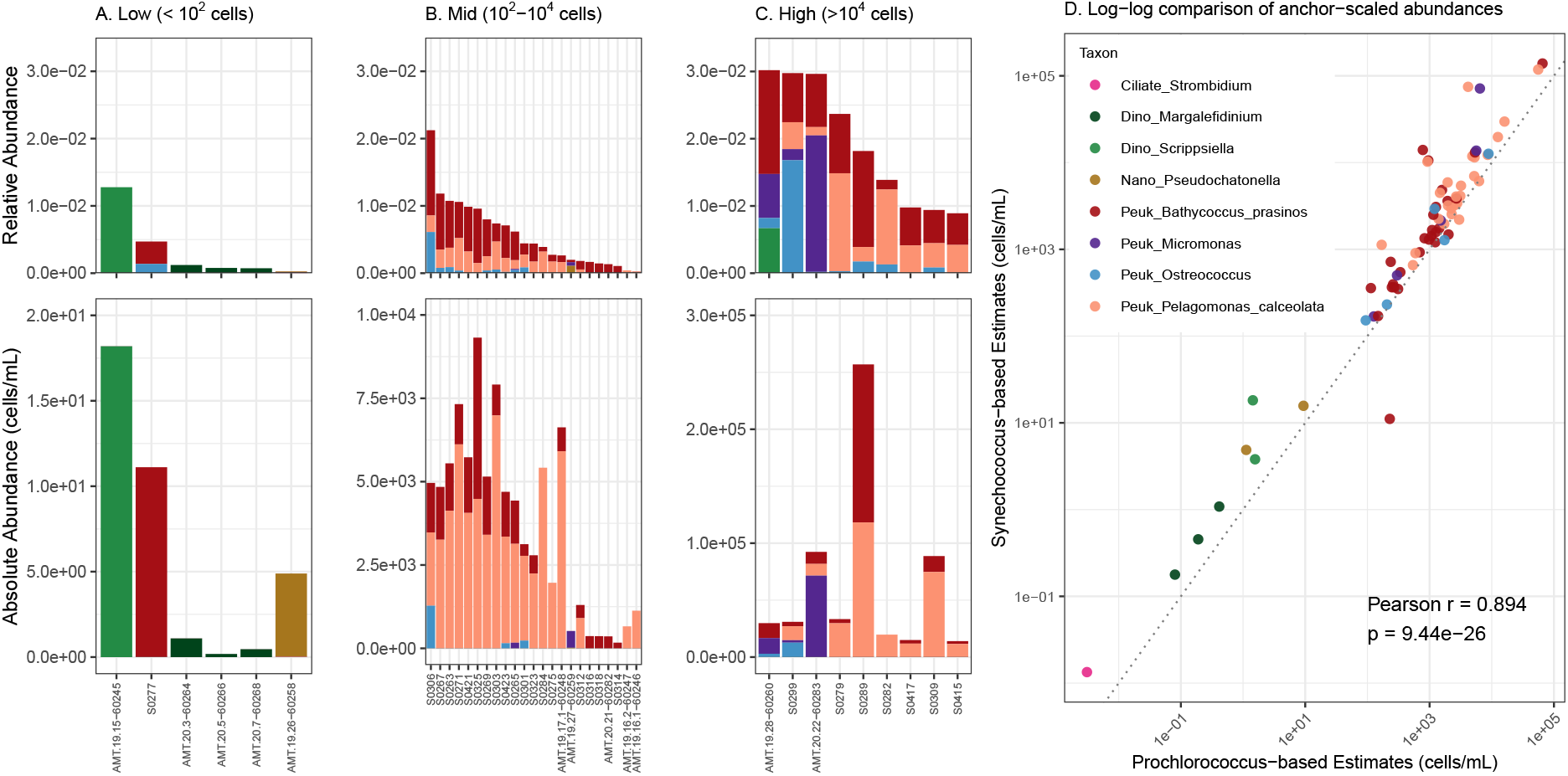
(A-C) *Synechococcus*-anchored cell concentration estimates for a variety of eukaryotic plankton for which reliable *ti* estimates were available. Samples across the AMT19, AMT20, and GP13 transects were stratified by total community cell abundance: (A) low group (*<* 10^2^ cells sample*^−^*^1^), (B) medium group (10^2^–10^4^ cells sample*^−^*^1^), and (C) high group (*>* 10^4^ cells sample*^−^*^1^). (D) Log–log comparison of *Synechococcus*-anchored versus *Prochlorococcus*-anchored cell concentration estimates across all samples, showing consistency between anchoring approaches and a systematic offset toward higher *Synechococcus*-based estimates across a wide range of taxa, including picoeukaryotes, nanoplankton, dinoflagellates, and ciliates.

Furthermore, to evaluate potential systematic biases introduced by the choice of anchor taxon, we compared absolute gene and cell concentrations estimated for non-cyanobacterial taxa using *Synechococcus* versus *Prochlorococcus* as anchors. Across all three transects, cell concentration estimates based on the *Synechococcus* anchor were generally higher than those based on *Prochlorococcus* (Fig. 6), consistent with the pattern observed for the cross-anchored picocyanobacteria (Fig. 4). Similarly, the *Syn*-anchored gene concentration estimates were systematically higher than *Pro*-anchored estimates for a wide range of prokaryotic and eukaryotic taxa within the top 15 ASVs along the AMT19 transect (Supplementary Figs. S2, S3). This reinforces that while ASV-calibration recovers quantitative abundances for diverse taxa, the choice of anchor systematically shifts absolute estimates, highlighting the diagnostic value of cross-anchoring in identifying and accounting for potential biases.

### 3.4 Evaluating ecotype-level cell concentrations and biogeography

Lastly, we applied the ASV-calibration framework to examine the distribution of *Prochlorococcus* ecotypes along the GP13 transect where we had metabarcode and flow cytometry samples (∼153–190°E). Temperatures in the upper 30m of the euphotic zone ranged from ∼19.3 to 23.5°C and exhibited a clear longitudinal temperature gradient, with warmer western waters (relative to cooler eastern waters). Correspondingly, absolute cell abundances of *Prochlorococcus* ecotypes (assuming constant *t_i_* = 1 across ecotypes) inferred from ASV data calibrated with *Synechococcus* as the anchor spanned wide ranges and revealed a distinct ecotype transition along this temperature gradient.

The combination of the deep taxonomic resolution of metabarcodes along with quantitative calibration, affords a view of absolute population concentrations at the ecotype level along the transect. Across the full section of the transect, *Pro*-HLI and *Pro*-HLII were the most dominant ecotypes. Ambiguous *Prochlorococcus* ASVs, that did not correspond uniquely to HLII and HLI ecotypes occurred at lower but variable abundances across the transect. When the transect was partitioned at ∼175°E to reflect the longitudinal temperature gradient in the upper 30m, distinct differences in ecotype distributions were evident (Fig. 7). In the warmer western section (*<* 175°E), *Pro*-HLII was prevalent, with *Pro*-HLI populations orders of magnitude lower . In contrast, *Pro*-HLI increased substantially in the cooler eastern section (≥ 175°E) while *Pro*-HLII remained abundant but at lower typical values than in the west (Supplemental Table 1). The ecotype-level taxonomic resolution of *Prochlorococcus* afforded by the GRUMP data by itself only results in relative abundances, whereas the flow cytometry cell counts of *Prochlorococcus* provide absolute abundances but cannot yield ecotype level distinctions. By combining the two we leverage the strengths of each independent methodology to generate ecotype-level cell count estimates.

**Figure 7:**
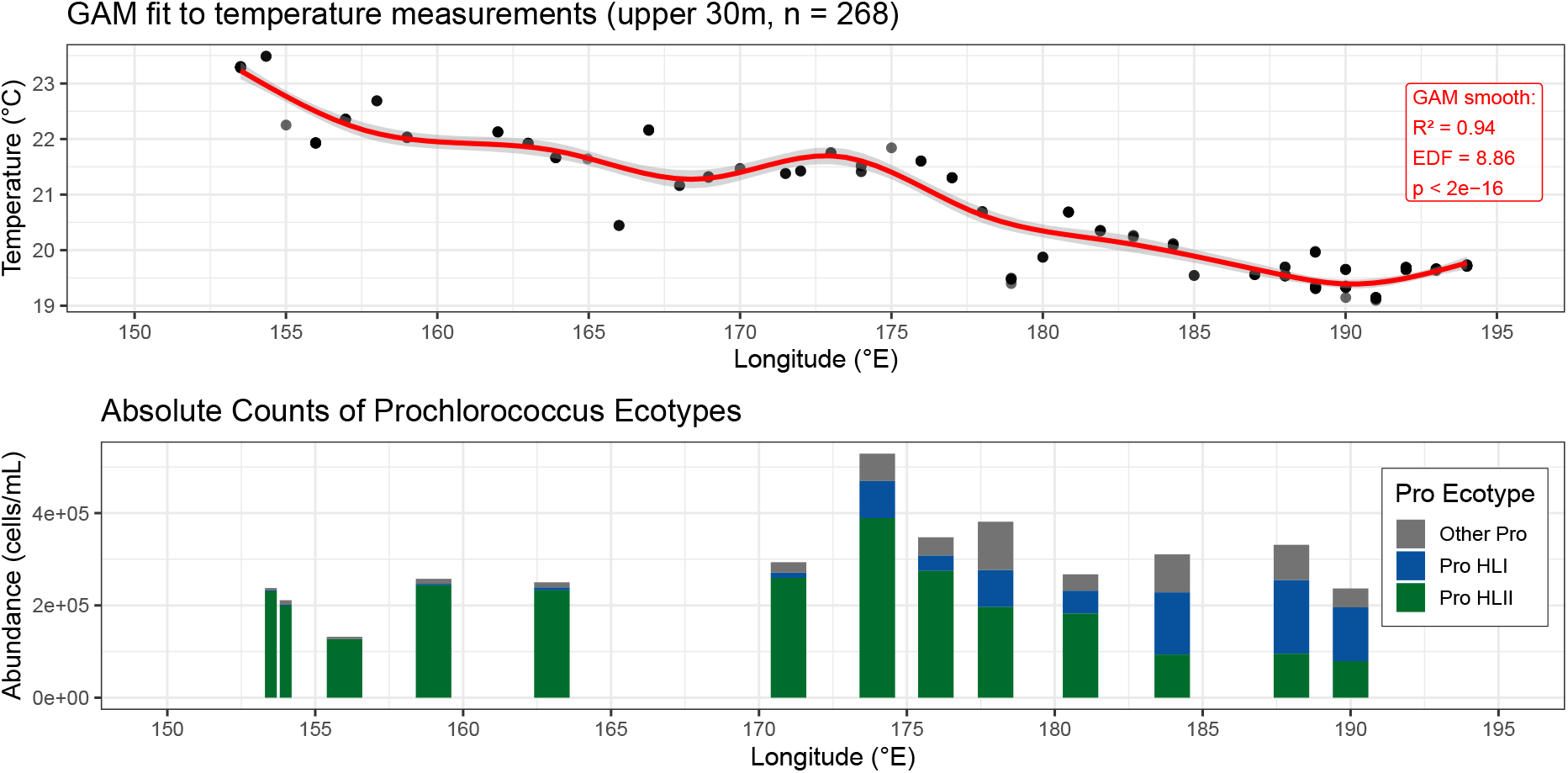
(a) Generalized Additive Model (GAM) fit to temperature measurements in the upper 30m of the water column (n = 268) along the first leg of the GP13 transect (longitude 150–195°E), showing warmer western waters (*<* 175°E) and cooler eastern waters (*>* 175°E). (b) Absolute abundances of *Prochlorococcus* ecotypes HLII and HLI sampled within the upper 30m of the euphotic zone, show a transition from HLII (green) dominance in the west to HLI (blue) dominance in the east along the GP13 temperature gradient.

## 4 Discussion

Quantitative molecular characterization of microbial abundances is opening the door for efficient and systematic mapping and monitoring of taxonomically-resolved biogeography of the ocean and other environments. Traditionally, such characterizations of marine plankton, derived from metabarcoding or metagenomics, have provided only compositional data as relative abundances in gene currencies (Sunagawa et al., 2015; McNichol et al., 2025). While internal nucleic acid standards (“spike-in”) (Lin et al., 2019; Satinsky et al., 2013) have occasionally been used to provide absolute gene concentrations (Jones-Kellett et al., 2024; Bei et al., 2025), the method remains underutilized in marine microbial ecology. By cross-calibration with flow cytometer (or other cell counting methods) the full potential of many existing molecular datasets could be extended to afford cross-sample comparisons and quantitative ecological inference across studies.

Building on previous work along the G4 transect, which demonstrated a tight correspondence between picocyanobacterial cell counts derived from internal standards and flow cytometry (Jones-Kellett et al., 2024), our framework successfully recapitulates these relationships for both *Prochlorococcus* and *Synechococcus* using calibrated ASV-based cell abundances (Figure 2). The study further demonstrates that (1) anchor choice serves as a diagnostic tool for assessing data reliability and potential biases, (2) estimating absolute gene concentration shifts interpretation of compositional data, and (3) further calibration to absolute cell counts expands our ecological interpretation. Below, we discuss the main findings, their implications, and remaining uncertainties.

### 4.1 Anchor Choice as a Diagnostic Tool

Choosing an appropriate anchor taxon is central to the calibration framework and provides a built-in diagnostic for assessing the reliability of absolute abundance estimates. The method described here is robust to a number of normal variations across study design and data characteristics. The sample-specific correction factor *a_j_* (Figure 3) is simply a way to transform compositional data into absolute data using the anchor taxon so high sequencing depth is not a significant requirement as long as the taxa of interest are detectable.

Secondly, this technique does not rely on a universal primer; the approach could just as easily be applied to a 16S-only, 18S-only, or group-specific primer sets. Finally, it is not dependent on PCR amplicon sequencing. So long it is possible to align taxonomic units (i.e., OTUs/ASVs) with independently-measured cell counts for an environmental taxon, it will be possible to use this framework. For example, microscopic counts of a eukaryotic phytoplankter or CARD-FISH counts of a bacterioplakton ((Moter and Gobel, 2000; Amann et al., 1995)) could be used as an anchor taxon for an 18S-specific or 16S-specific amplicon dataset, respectively. Similarly, metagenomically-derived SSU rRNA taxa or OTUs could be used so long as it is possible to accurately link those data with the anchor taxon.

We found that cross-anchoring between different taxa further enhances the diagnostic power of the approach. Across multiple transects (AMT-19, AMT-20, GP13), cross-anchoring between *Prochlorococcus* and *Synechococcus* preserves relative abundance patterns but produces systematic offsets in absolute cell concentrations (Figure 4). The offset in this case is likely due to the challenges of detecting *Prochlorococcus* in high light conditions, suggesting that (in this particular instance) *Synechococccus* may be the more reliable anchor. However, in other situation low abundances of *Synechococcus* might make it a less reliable anchor. Cross anchoring provides some important checks on the calibration procedure and some quantification on systematic uncertainties. We note that when the calibration is applied across a wide taxonomic range, the correspondingly wide range of cell concentrations is captured faithfully and the offsets become small relative to the range in absolute cell concentration (Fig. 7).

### 4.2 Absolute Gene Concentration

Transforming relative sequence abundances into absolute gene concentrations provides a fundamentally more informative metric for comparing microbial populations across samples, transects, or studies. By extending absolute abundance estimation to the 15 most abundant ASVs along the AMT-19 transect, we revealed community and individual level patterns obscured in relative abundance data (Fig. 5, Supplementary Fig. S2). Absolute gene concentrations provide an integrated view of microbial biogeography and enable researchers to quantify real changes in total gene content. However, interpretation of taxa with unknown or highly variable genome copy numbers (*t_i_*) remain challenging, and thus, estimates for larger eukaryotic plankton should be interpreted cautiously. Anchor choice also continues to influence absolute gene concentration estimates, emphasizing the need for careful calibration (Supplementary Fig. S3), while demonstrating the added value of uncertainty bounds acquired by leveraging the cross-anchoring approach (Supplementary Fig. S2).

Several potential limitations and biases remain when inferring absolute gene concentrations from metabarcoding data derived from complex environmental mixtures. Biases can arise at multiple stages, including DNA extraction, PCR amplification, sequencing, and bioinformatic processing, but many are correctable using one or more calibration standards composed of environmental samples (or mixtures of samples) that cover all taxa present in a given environment (McLaren et al., 2019). Additionally, this approach requires expert knowledge of the system, knowledge of gene copy numbers per cell for at least one “anchor taxon” present across all samples, and appropriate annotation of metabarcoding data. In oceanic environments, the rapid accumulation of genomic information (Biller et al., 2018b), improvement of environmental databases (Quast et al., 2012; Larkin et al., 2021a), and ongoing efforts to intercalibrate rRNA copy numbers for ecologically-relevant organisms make these challenges increasingly tractable. In the future, intercalibration of data derived from this technique with independent, expert-derived measurements of cell numbers or gene copy numbers from internal standards could provide the clearest indication of whether this technique is broadly useful in marine systems and how it compares with other high-throughput methods of community characterization such as flow cytometry (Swalwell et al., 2011b), IFCB (Olson and Sosik, 2007b), or UVP (Picheral et al., 2022).

### 4.3 Beyond Gene Concentrations: Conversion to Cell Concentrations

It is interesting to consider the potential to calibrate absolute gene concentrations to cell concentration, the intrinsic currency of cell counting approaches: for example, merged data sets obtained using different technologies could be used to improve biogeographical coverage. Extending calibration further to estimate biomass, which can often be derived from light-scattering measurements by flow-cytometry (e.g., Ribalet et al. (2019)) would provide opportunities to inform carbon and nutrient budget studies and provide a direct interface to ecosystem and climate models. While challenging, this goal is potentially very valuable for biogeochemical interpretations, where mass budgets become of particular interest.

Applying the ASV-calibration framework to *Prochlorococcus* ecotypes along the GP13 transect demonstrates the ability to resolve ecotype-specific abundance patterns across a longitudinal temperature gradient in the western Pacific. Warmer western waters favored the dominance of HLII ecotypes, whereas cooler eastern waters supported higher relative abundances of HLI ecotypes (Fig. 7). These observations are consistent with prior studies linking *Prochlorococcus* ecotype distributions across temperature regimes in laboratory and in situ conditions (Johnson et al., 2006; Biller et al., 2015), and illustrate that absolute cell counts derived from calibrated ASVs can capture ecologically meaningful patterns of temperature-driven niche partitioning along environmental gradients. Importantly, this approach combines the taxonomic resolution of metabarcoding with quantitative scaling, providing ecotype-level cell abundances that neither relative sequence data nor bulk flow cytometry alone can deliver.

While current *t_i_*knowledge is well documented for the prokaryotic subset of microbial taxa such as *Prochlorococcus* or *Synechococcus* (Dufresne et al., 2003; Fuller et al., 2003; Schirrmeister et al., 2012; Stoddard et al., 2015), it remains much more difficult to characterize for eukaryotic plankton, which can have much larger variability in rRNA gene copy numbers (Martin et al., 2022). Nevertheless, for a subset of eukaryotic ASVs present across the AMT-19, AMT-20, and GP13 transects with well-characterized *t_i_* values (Martin et al., 2022), converting gene concentrations into cell concentrations revealed pronounced deviations from relative abundance metrics. Larger organisms such as nanophytoplankton and dinoflagellates, which carry higher rRNA copy numbers, often appear overrepresented in relative abundance data compared to picoeukaryotic plankton. When calibrated to cell counts, these discrepancies are corrected, showing that actual cell abundances for larger taxa are often orders of magnitude lower than relative sequence abundances alone would suggest (Fig. 6). Across all three transects, estimates based on the *Synechococcus* anchor were also generally higher than those based on *Prochlorococcus* (Fig. 6), consistent with the pattern observed in cross-anchored picocyanobacteria (Fig. 4). This underscores that while ASV-calibration provides quantitative abundances for diverse taxa, anchor choice systematically shifts absolute estimates, highlighting the diagnostic value of cross-anchoring to identify and account for potential biases.

Collectively, these results demonstrate that the ASV-calibration framework can be extended beyond cyanobacteria to provide quantitative, interpretable measures of plankton community structure. Even in the absence of *t_i_*, absolute gene concentrations capture variations obscured in relative abundance data, while conversion to cell concentrations for known taxa allows more direct ecological interpretation. Remaining limitations include uncertainty and incomplete knowledge of *t_i_* across diverse taxa, potential PCR and primer biases, and the influence of anchor taxon variability on propagated errors. Additionally, assumptions in converting gene concentration to cell concentration or biomass introduce further uncertainty. While the framework represents a significant improvement over purely compositional data, these uncertainties must be explicitly considered when interpreting absolute abundance estimates.

### 4.4 Concluding Remarks

Moving beyond relative abundances of planktonic taxa will be a valuable additional step for quantitative microbial ecology. Absolute abundance provides additional information and makes additional analyses possible, including aggregating samples and merging data sets in ways not possible with relative abundances. Relative abundances can, of course, always be derived from absolute. Calibrating relative gene abundances to absolute environmental gene concentrations is the first step towards calibration into cell concentration (with knowledge of gene copies per cell) and potentially even biomass (with additional knowledge of cell volumes and carbon concentrations). Though these additional requirements represent some significant challenges today, the potential for efficient, quantitative surveys of taxonomically resolved gene concentrations, cell concentrations, and biomass using efficient, calibrated molecular tools could transform quantitative studies of ocean biogeochemistry and marine microbial ecology.

## 5 Author contribution

E.S-G., M.F. and J.M. conceived the study. E.S-G. and Y.R. performed analysis and simulations. E.S-G., Y.R, M.F. and S.D. wrote the manuscript. E.S-G., Y.R., J.M., F.R., S.D., J.F. and M.F. commented on the results and edited the manuscript. G.T. and C.H provided the data and commented on the results.

## 6 Supplementary Figures

**Supplementary Figure S1:**
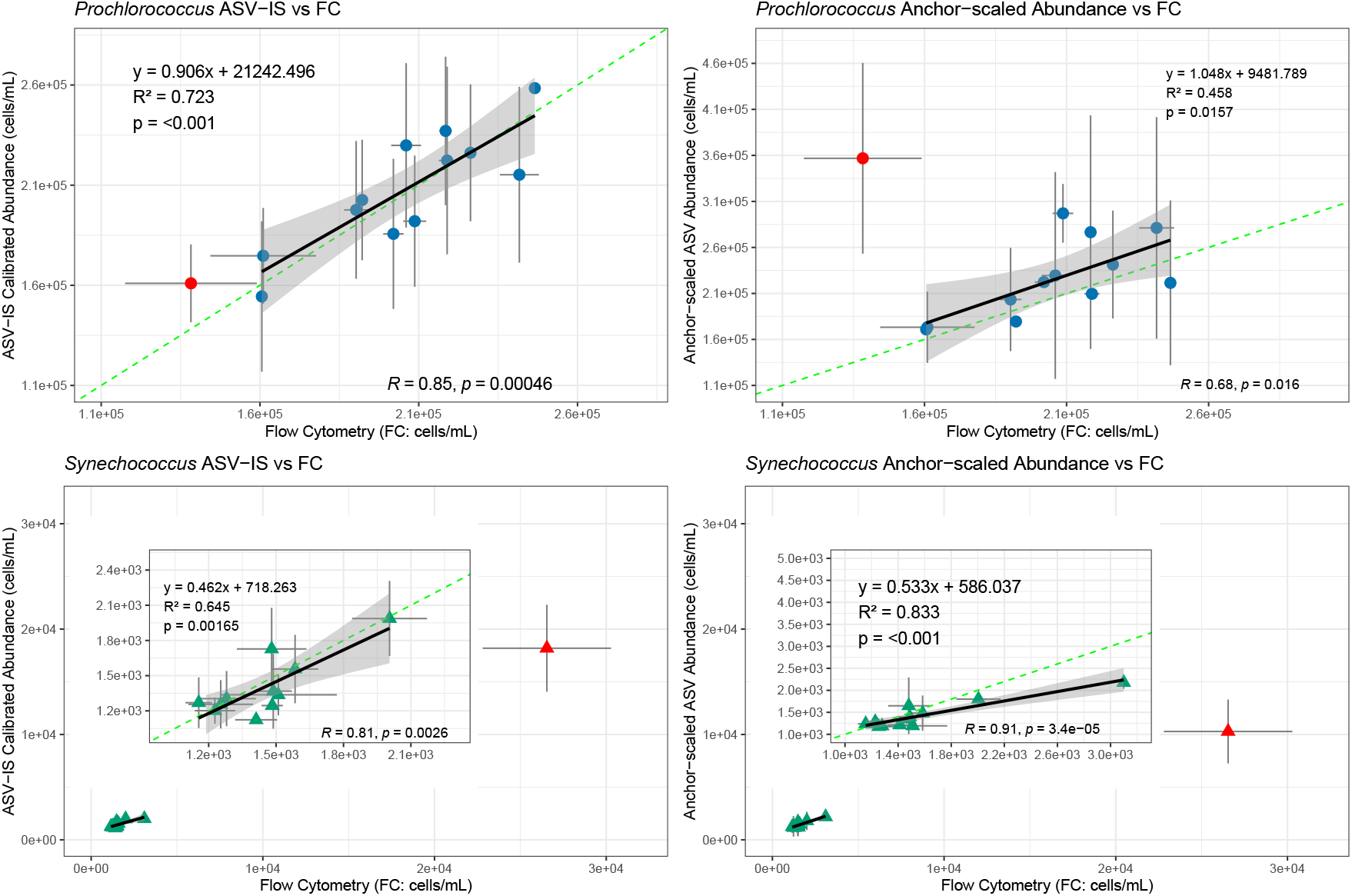
Comparison of ASV-based and flow cytometry–based cell concentrations for *Prochlorococcus* and *Synechococcus*. ASV-internal standard (ASV-IS) calibrated absolute abundances versus flow cytometry (FC) counts for *Prochlorococcus* (top left) and *Synechococcus* (bottom left) (Jones-Kellett et al., 2024). Anchor–scaled abundance for *Prochlorococcus* (derived using *Synechococcus* as anchor) versus FC counts (top right) and for *Synechococcus* (derived using *Prochlorococcus* as anchor: bottom right). Points represent mean cell abundances (cells mL*^−^*^1^) across replicates, with horizontal and vertical error bars indicating ±1 SD. Linear regressions (solid black lines) and corresponding 95% confidence intervals were fitted excluding sample 10 (highlighted in red). The dashed green line represents the 1:1 relationship. Insets in *Synechococcus* panels show zoomed views of the lower abundance range. Regression equations, coefficients of determination (*R*^2^), and Pearson correlation statistics are displayed within each panel.

**Supplementary Figure S2:**
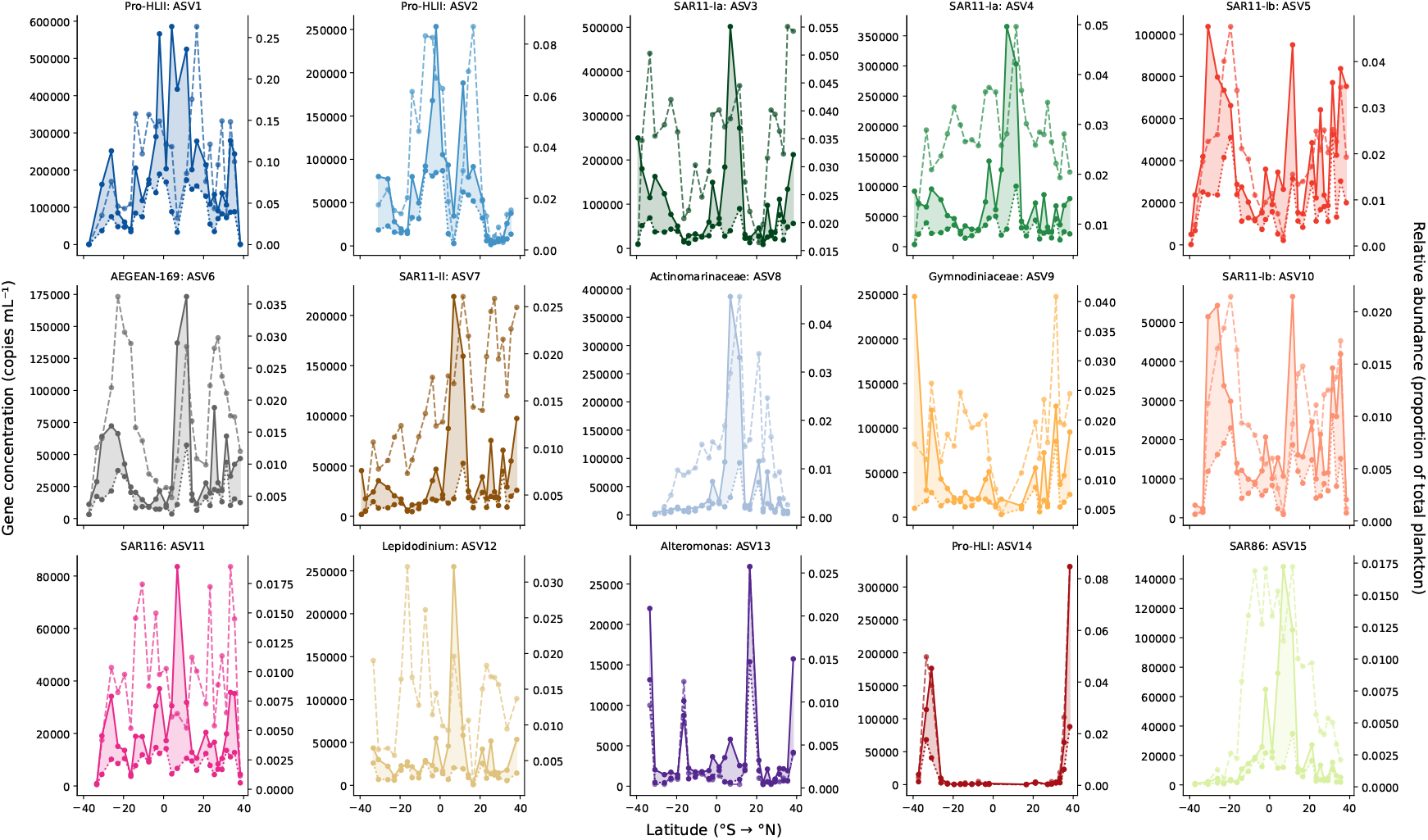
Latitudinal distribution of the top 15 ASVs along the AMT19 transect shown as relative abundance (proportion of the total plankton community; right y-axis; dashed line) and absolute gene concentration estimate (left y-axis) that is anchored by *Prochlorococcus* (dotted line) and *Synechococcus* (solid line). The shaded region between the dotted and solid line corresponds to the bound between the upper and lower gene concentration estimates provided by the cross-anchoring approach.

**Supplementary Figure S3:**
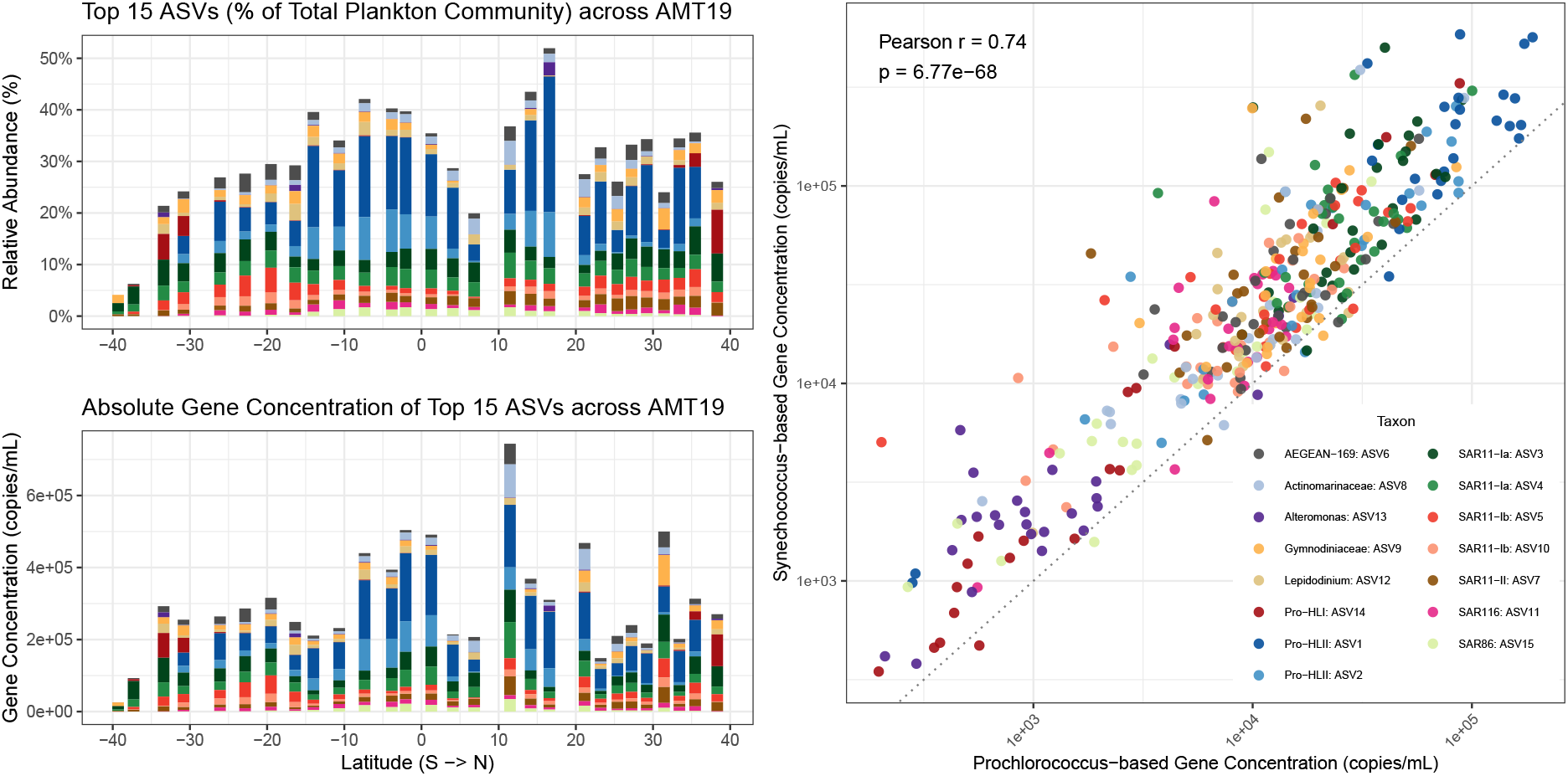
A) Latitudinal distribution of the top 15 ASVs along the AMT19 transect shown as relative abundance (of the total plankton community; top panel) and absolute gene concentration anchored using *Prochlorococcus* as the alternate anchor (bottom panel). B) Scatterplot showing the relationship between *Syn*-anchored versus *Pro*-anchored estimates of gene concentration of the top 15 ASVs along the AMT19 transect.

**Supplementary Figure S4:**
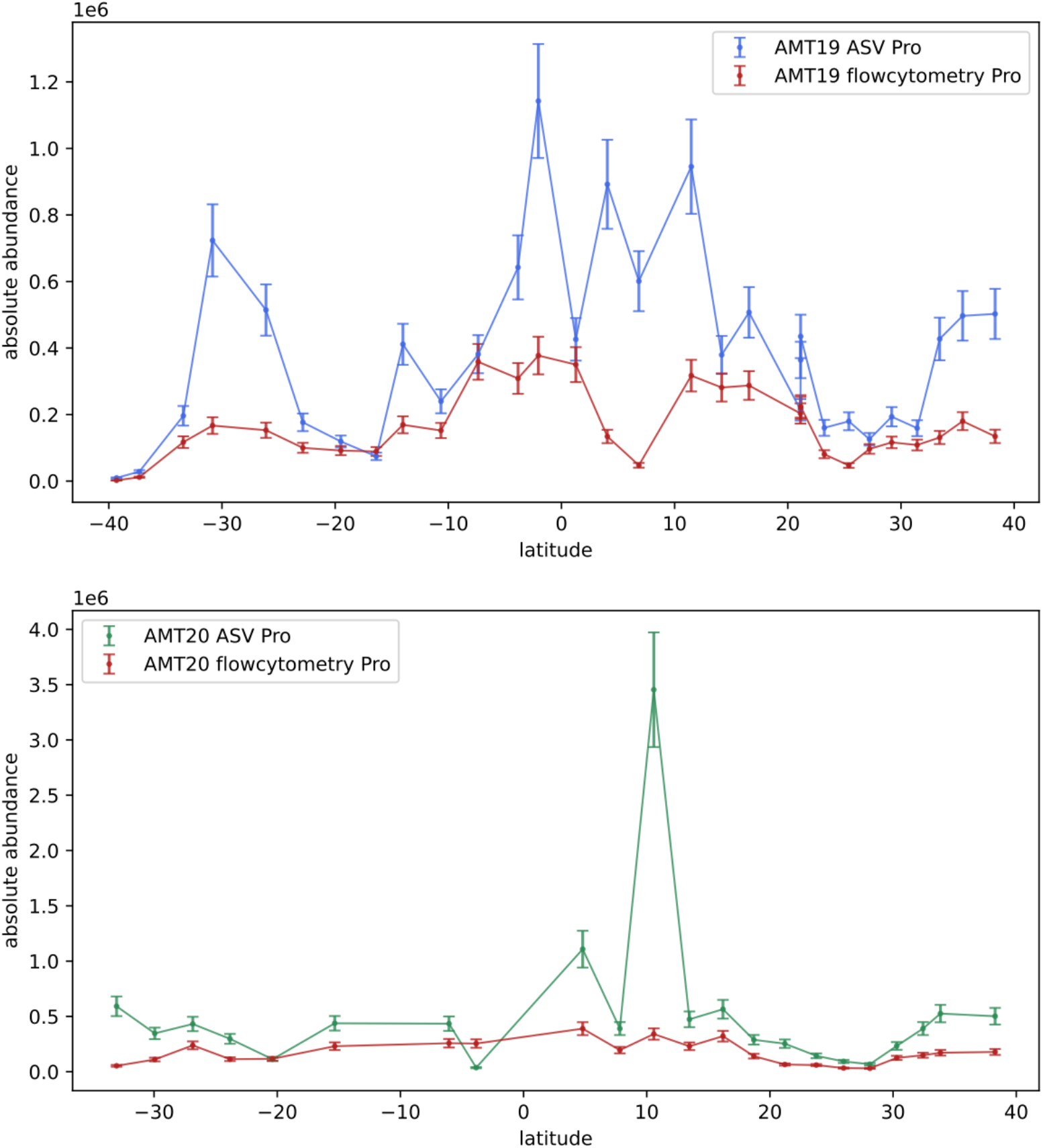
Flow cytometric and calibrated metabarcode absolute cell concentration of *Prochlorococcus* along AMT19 (a) and AMT20 (b). ASV values calibrated using *Synechococcus* anchor.

**Supplementary Figure S5:**
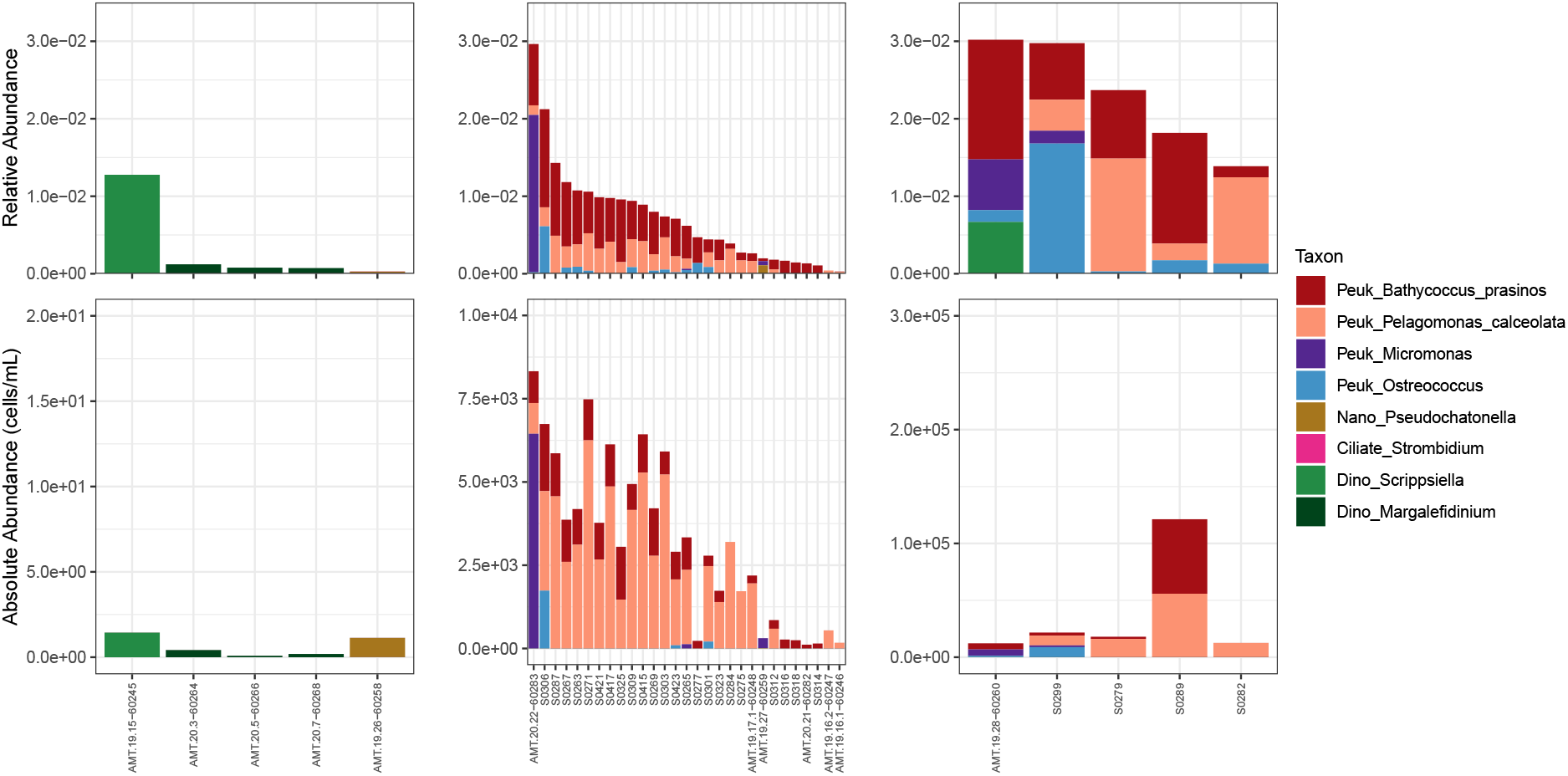
*Prochlorococcus* anchored cell concentration estimates for eukaryotic plankton for which reliable *ti* estimates were available.

**Table S1:** *Prochlorococcus* abundances and associated errors across samples.

| SampleID | HLI_abs | HLI_err | HLII_abs | HLII_err | Other_abs | Other_err |
| --- | --- | --- | --- | --- | --- | --- |
| S0263 | 3234 | 840.32 | 243525 | 63269.61 | 10762 | 2796.12 |
| S0269 | 6480 | 1683.63 | 232555 | 60419.49 | 11034 | 2866.72 |
| S0279 | 11521 | 2993.28 | 259614 | 67449.60 | 22445 | 5831.47 |
| S0284 | 134859 | 35037.52 | 93475 | 24285.43 | 82231 | 21364.34 |
| S0292 | 80516 | 20918.56 | 389069 | 101083.14 | 59224 | 15386.76 |
| S0299 | 1922 | 499.42 | 230549 | 59898.32 | 5182 | 1346.27 |
| S0301 | 1866 | 484.76 | 199432 | 51813.86 | 9876 | 2565.97 |
| S0306 | 0 | 0.00 | 127064 | 33012.24 | 4845 | 1258.76 |
| S0312 | 33544 | 8714.94 | 274733 | 71377.76 | 39324 | 10216.56 |
| S0316 | 116593 | 30291.78 | 79251 | 20590.08 | 40865 | 10616.96 |
| S0323 | 159742 | 41502.31 | 95273 | 24752.53 | 76224 | 19803.56 |
| S0415 | 80293 | 20860.80 | 196407 | 51027.97 | 104589 | 27173.03 |
| S0421 | 49461 | 12850.47 | 182504 | 47415.99 | 35347 | 9183.46 |

